# Molecular epidemiology of human respiratory syncytial virus with severe acute respiratory infection in Huzhou from 2016 to 2019

**DOI:** 10.1101/2021.02.15.431241

**Authors:** Deshun Xu, Lei Ji, Xiaofang Wu, Wei Yan, Liping Chen

**Author notes:** Corresponding Author: Liping Chen, Huzhou Center for Disease Control and Prevention, 999 Changxing Road, Huzhou, Zhejiang 313000, China.

## Abstract

**Background:** Human respiratory syncytial virus (HRSV) is one of the major cause of acute lower respiratory infection in infants, the elderly and people with low immunity worldwide. Based on antigenic and genetic variations, Human respiratory syncytial virus is divided into two subgroups (A and B). Each of the subgroups is further categorized into genotypes based on the phylogenetic analyses of the sequences of the second hypervariable region.

**Methods:** Nasopharyngeal swabs (NPSs) were collected from patients of the First People’s Hospital in Huzhou from January 2016 to December 2019. Real-time RT-PCR (qPCR) was performed using double nucleic acid detection kit for respiratory syncytial virus (A\B) (Shenzhen shengkeyuan) with the ABI Q7 (Applied Biosystems). For genotyping, the primer set A-F/A-R was used to amplify the G protein of HRSV-A. Primer set B-F/B-R was used to amplify the G protein of HRSV-B. The phylogenetic analysis was constructed using the neighbor-joining algorithm with the Kimura two-parameter model and supported statistically by bootstrapping with 1000 replicates with MEGA software (version 7.0) with 1000 bootstrap replicates.

**Results:** A total of 973 nasopharyngeal swab samples were collected from January 2016 to December 2019, and 63 samples were positive for RSV nucleic acid, with the detection rate of 6.47%. Of the positive specimens, 28 were belonged to HRSV-A, and 35 were belonged to HRSV-B. Infection with RSV was found in all age groups tested, with the 0-1 year age group having the highest detection rate 15.2%. The detection rate was high from November to next March. Phylogenetic analysis clustered HRSV-A strains identified in Huzhou into ON1genotype. All 17 of the HRSV-B strains belonged to BA9 genotype.

**Conclusions:** We analyzed the HRSV strains circulation in Huzhou from January 2016 to December 2019 in Huzhou, China. This is the first molecular analysis on HRSV in Huzhou. We found Subgroup A and B of RSV were co-circulating and the 0-1 year age group having the highest infection rate.

## Background

Human respiratory syncytial virus (HRSV) is one of the major causes of acute lower respiratory infection in infants, the elderly and people with low immunity worldwide [1]. In the first year of life, nearly all children are infected once or multiple times with HRSV [2]. The virus is known to cause repeated infection throughout life [3–4].

Human respiratory syncytial virus was classified in the genus Orthopneumovirus of the Pneumoviridae family within the order Mononegavirales [5]. HRSV is an enveloped virus with single-stranded negative RNA of about 15.2 kb in length. The genome encodes 11 proteins, NS1, NS2, N, P, M, SH, G, F, M2-1, M2-2 and L. HRSV is divided into two subgroups (A and B), based on antigenic and genetic variations [6]. Glycoprotein G gene is the most variable among HRSV genes and has two hypervariable regions. The analysis of this region is commonly used for studying the molecular epidemiology studies of HRSV [7]. According to the different nucleotide sequences of the second hypervariable region of G protein gene, the subtypes can be further divided into several genotypes [8].

Here, we describe the prevalence and genetic characteristics of HRSV from January 2016 to December in 2019 in Huzhou City, Zhejiang, China, and according to the G protein gene we analysised the subtypes circulating in Huzhou.

## Methods

### Specimen collection

This study was conducted at the First People’s Hospital in Huzhou, a national sentinel hospital for monitoring cases of severe acute respiratory infection. Samples used for this study were collected from January 2016 to December 2019. Nasopharyngeal swabs (NPSs) from patients who attend the hospital with severe acute respiratory infection were put into the same vial of virus transport medium. The samples were transported on ice to the microbiology laboratory of Huzhou Center for Disease Control and prevention for immediate storage at −70 °C prior to analysis.

### HRSV detection/Real-time PCR for subgrouping

Viral RNA is extracted from 200μl of the clinical specimens using a QIAamp viral RNA mini kit (Qiagen, Hilden, Germany) according to the manufacturer’s instructions.The RNA extracts were subjected directly to reverse transcription (RT)-PCR (Polymerase Chain Reaction) or stored at −70 °C. Real-time RT-PCR (qPCR) was performed using double nucleic acid detection kit for respiratory syncytial virus (A/B) (Shenzhen shengkeyuan) with the ABI Q7 (Applied Biosystems). The reaction was conducted according to the manufacturer’s instructions with total volume of 20 μL.

### DNA sequencing of G protein gene of RSV

For genotyping, the primer set A-F: TATTCATATCATCGTGCTTATACAAGTTA/A-R: GCAGGGTACAAAATTGAACACTT was used to amplify the G protein of HRSV-A. Primer set B-F: GAAGTGTTCAACTTTGTTCCCTGTA/B-R: GGATGGTTGAGTAGAGAGATTGCT was used to amplify the G protein of HRSV-B. RT-PCR was carried out using the One Step RNA PCR Kit (TaKaRa Biotechnology Dalian, China, CAT: DRR057A) with the amplification conditions as follows: 50°C for 30 min, 95°C for 3 min, followed by 35 cycles of 94°C for 30sec, 58°C for 30 sec, and 72°C for 1.5 min, and a final step at 72°C for 7 min. After amplification, 5 μL of the PCR products was visualized by agarose gel electrophoresis. The residual PCR products were purified using a QIAquick PCR purification kit (Qiagen, Leusden, The Netherlands), and the purified products were sequenced directly at both ends with amplification primers by TaKaRa Biotechnology (Dalian, China).

### Sequence analysis and phylogenetic analysis

The RSV sequences of both the group A and B were downloaded from Genbank. The phylogenetic analysis was constructed using the neighbor-joining algorithm with the Kimura two-parameter model and supported statistically by bootstrapping with 1000 replicates with MEGA software (version 7.0) with 1000 bootstrap replicates [9].

### Nucleotide sequence accession numbers

The GenBank accession numbers for sequences obtained in this study are MW282960-MW282987.

## Results

### Epidemic characteristics of HRSV

A total of 973 nasopharyngeal swab samples were collected from January 2016 to December 2019, and 63 samples were positive for HRSV, with the detection rate of 6.47%. Of the 63 HRSV-positive specimens, 28 were belonged to HRSV-A, and 35 were belonged to HRSV-B. The detection rate was 2.88% (28/973) and 3.6% (35/973) for subtype A and subtype B respectively. From 2016 to 2019, the detection rate of HRSV is similar every year. There was no significant difference in HRSV detection rate between different years (χ^2^ = 0.617, P = 0.892). The detection rate was 7.28% (41/563) for male and 5.37% (22/410) for female. There was no significant difference between genders (χ^2^ = 1.439, P = 0.230) (Table1).

**Table 1.**
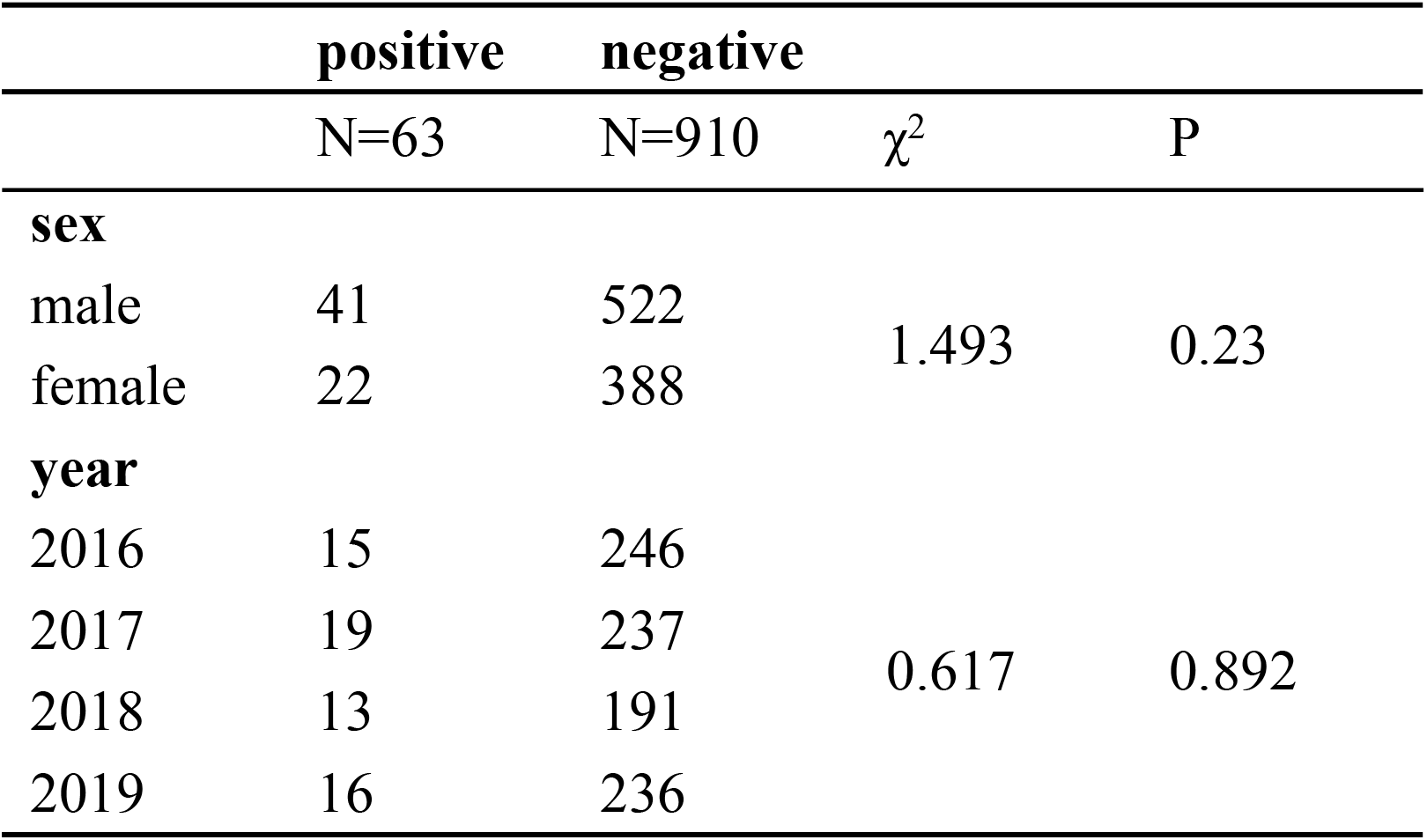
Epidemiological features of HRSV in Huzhou from January 2016 to December 2019

Infection with HRSV was found in all age groups tested, with the 0-1 year age group having the highest detection rate 15.2% (Fig 1). Then is the age group of 1-2 years old, with the detection rate of 12%. It is obvious that the detection rate decreases with the increase of age.

**Fig 1.**
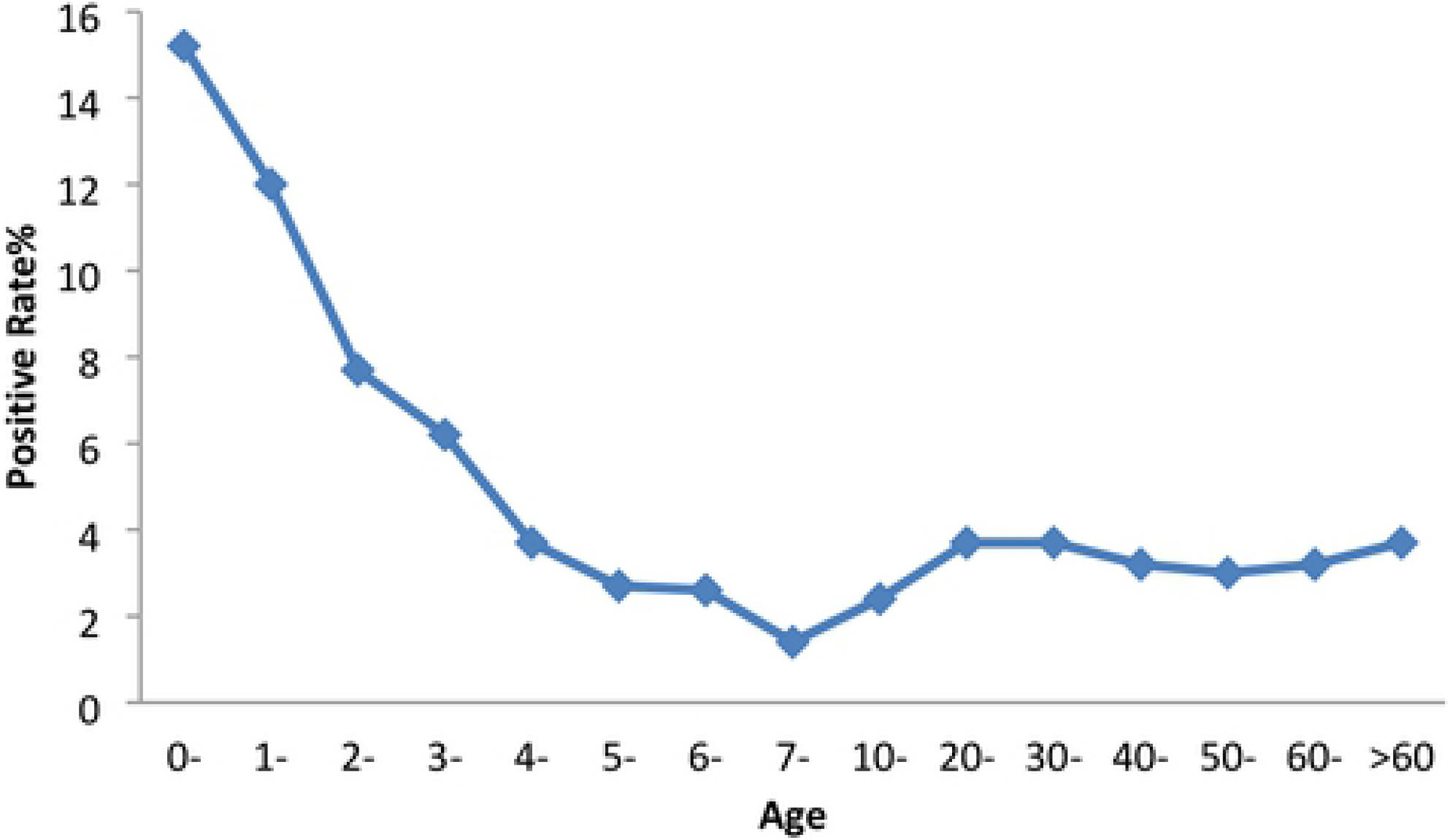
Age distribution of HRSV in Huzhou from January 2016 to December 2019. Age group distribution of HRSV in Huzhou from January 2016 to December 2019

Temporal distribution of HRSV infection was shown in Fig2. The HRSV detection rate increased from January, reached a peak in March, then declined and remains low between April and October. From November to December, HRSV detection rate increased suddenly and reached a peak in December. The detection rate was almost all over 10% from November to next March through the four years. In contrast, HRSV was not detected in some years from May to September.

**Fig 2.**
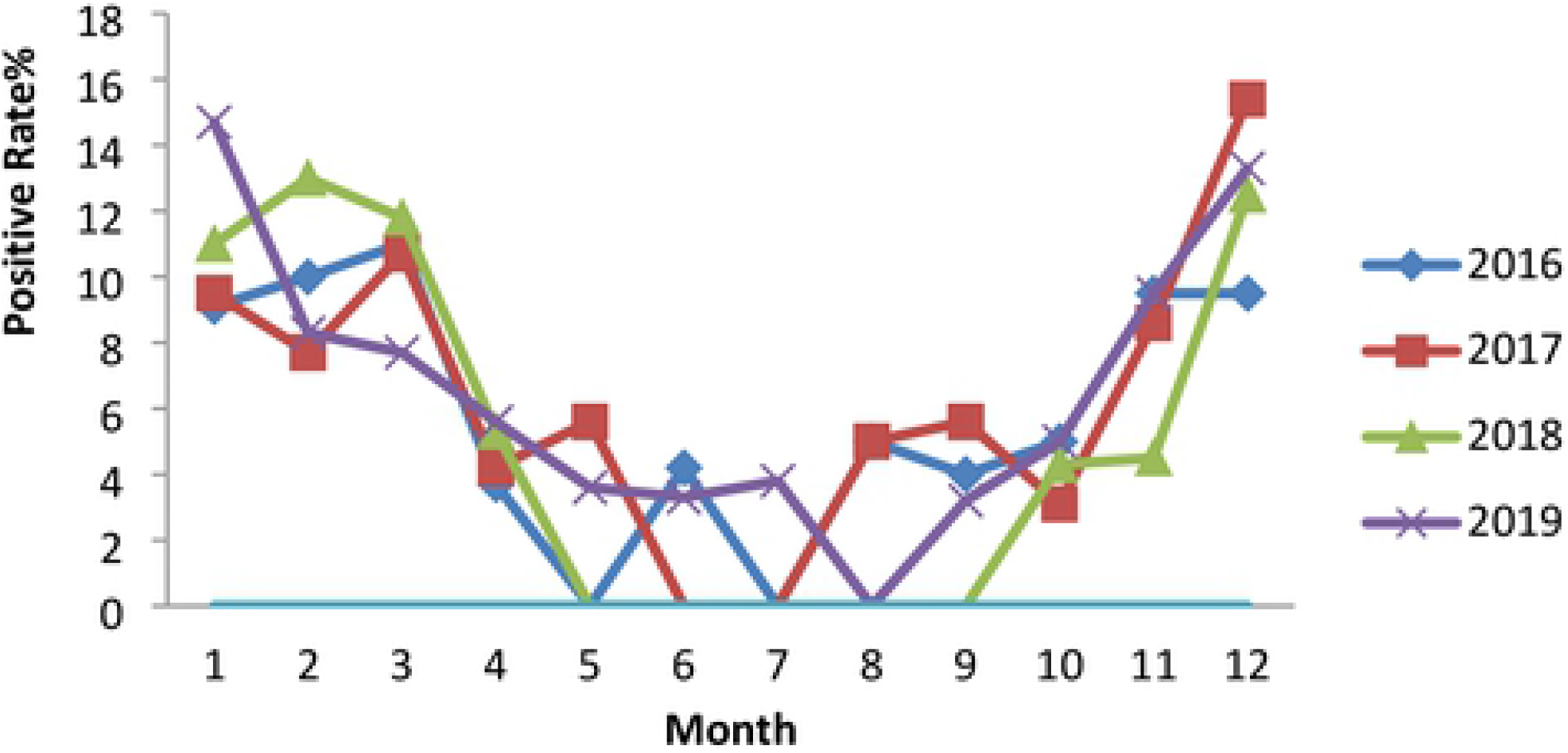
Temporal distribution of HARSV infection in Huzhou from January 2016 to December 2019. Monthly detection of HRSV in Huzhou from January 2016 to December 2019.

### Prevalence of HRSV genotypes

All samples that tested HRSV positive by real-time PCR were classified according to genotype. In total, 28 samples (44.4%) were successfully sequenced and genotyped by RT-PCR. Among them, 11 were HRSV-A and 17 were HRSV-B. The genetic sequences were not available for the other 17 HRSV-A and 18 HRSV-B which were real-time PCR positive. Phylogenetic trees were constructed on the basis of the G gene sequence by the neighbor-joining method. Phylogenetic analysis clustered HRSV-A strains identified in Huzhou into ON1 genotype, sharing the highest identity with the strain 2970-GD-CHN-2018 (MN007084) which isolated in Guangdong, China, 2018 and other three strains identified in Korea between 2014 to 2015. All 17 of the HRSV-B strains belonged to BA9 genotype and clustered with the strains which isolated in Korea, Japan, Philippines and Nicaragua from 2013 to 2015 (Fig 3).

**Fig 3.**
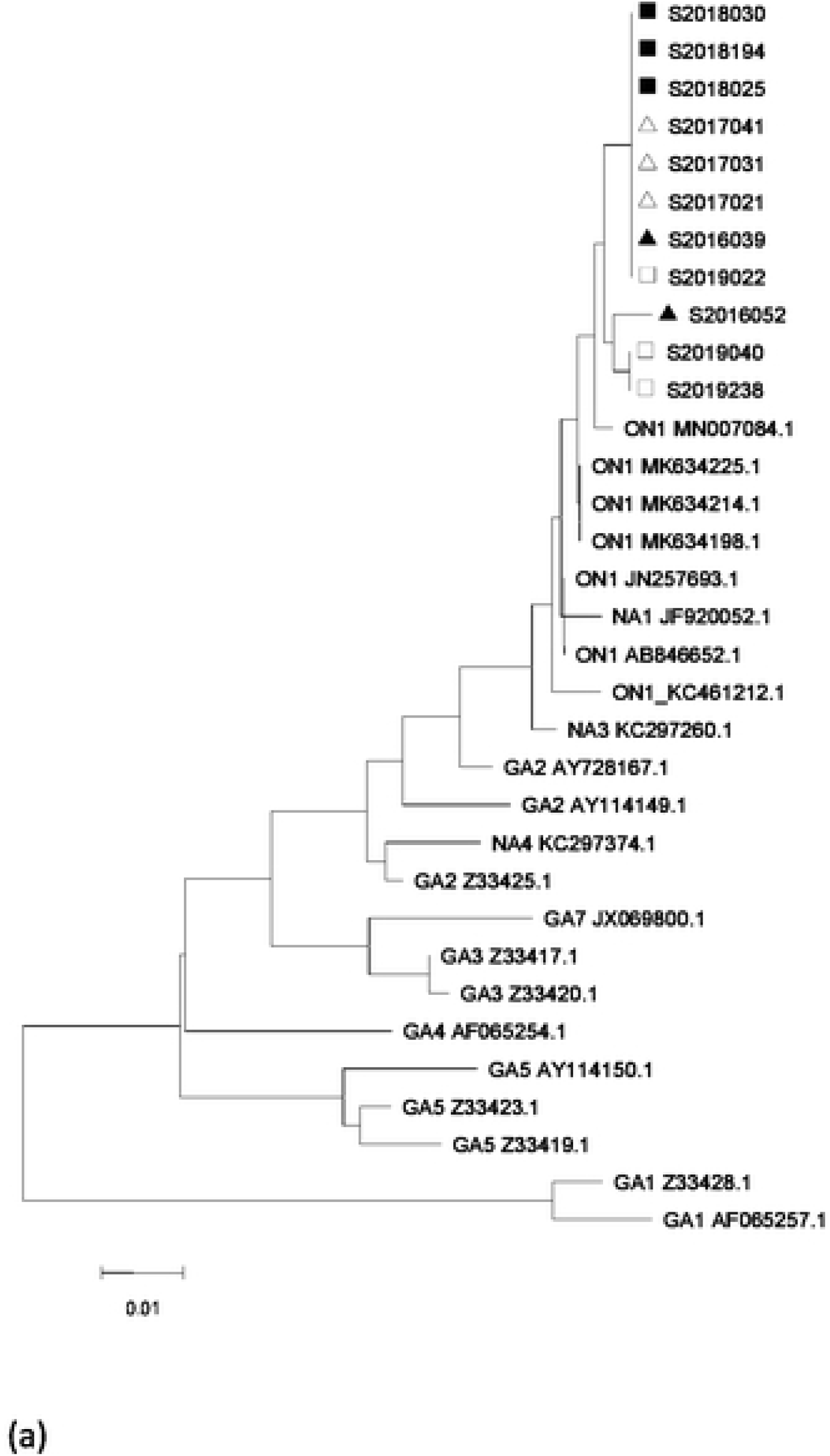
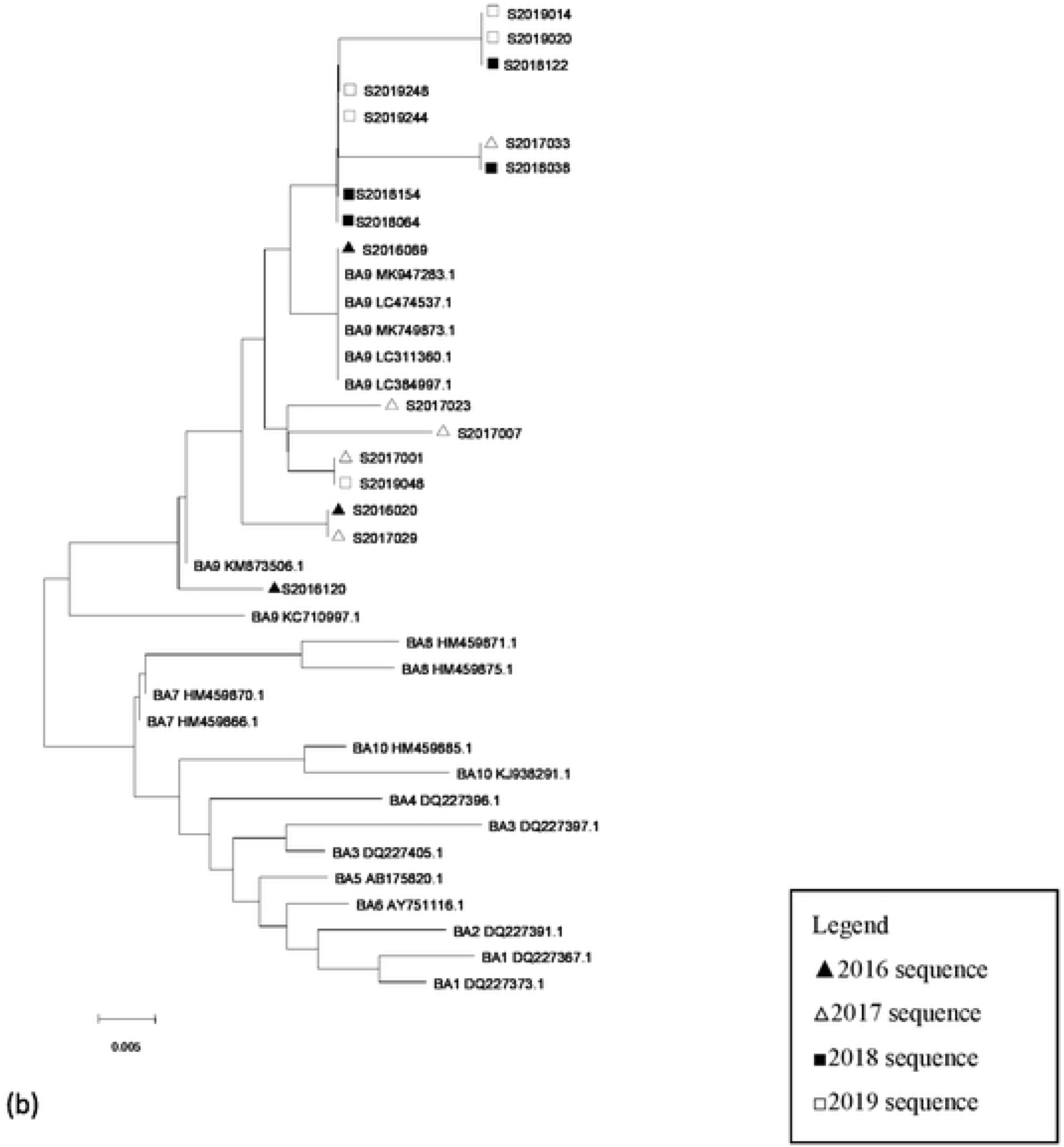
Phylogenetic analysis based on G gene sequence of HRSV-A (a) and HRSV-B (b). The trees were constructed by the neighbor-joining method with the Maximum Composite Likelihood model in MEGA (version 7.0), validated by 1000 bootstrap replicates. The samples are identified by data. Reference sequences (variants) are indicated by their genotypes and accession numbers. Phylogenetic analysis based on G gene sequence of HRSV-A (a) and HRSV-B (b) from severe acute respiratory infection patients in Huzhou from 2016 to 2019. The trees were constructed by the neighbor-joining method with the Maximum Composite Likelihood model in MEGA (version 7.0), validated by 1000 bootstrap replicates. The samples are identified by data. Reference sequences (variants) are indicated by their genotypes and accession numbers.

## Discussion

Human respiratory syncytial virus is a major cause of acute lower respiratory infection in infants and young children [10]. HRSV related acute lower respiratory infection brings a serious disease burden to human beings, especially in developing countries [11]. Adhesion protein G and fusion protein F are the main surface glycoproteins of HRSV, and they are the main antigens that stimulate the body to produce neutralizing antibodies. Although HRSV F protein is relatively conservative among subtypes or genotypes, it can induce high titer neutralizing antibody and has good cross protection effect. G protein also plays an important role in inducing and regulating immune response to infection, and the second hypervariable region of G protein is also a target gene for studying the molecular epidemiological characteristics of HRSV [12]. Recent studies have shown that two antibodies against G protein have been detected in adult B cells after HRSV infection, which can reduce lung infection and improve lung inflammation in mice [12,13].

Based on antigenic and genetic variations, Human respiratory syncytial virus is divided into two subgroups (A and B) [6]. The major variations between and within the two subgroups are found in the attachment glycoprotein G. Each of the subgroups is further categorized into genotypes based on the phylogenetic analyses of the sequences of the second hypervariable region. [14]

In this study, we investigated the genetic characteristics of circulating HRSV in Huzhou from January 2016 to December 2019. 63 were positive for HRSV among 973 samples, with 28 were belonged to HRSV-A, and 35 were belonged to HRSV-B. The detection rate was high from November to next March. Generally, the epidemic season of HRSV infection is in winter and spring every year. At the end of autumn, with the decrease of environmental temperature, the infection rate of HRSV gradually increased and reached the peak from December to next march [15]. In the subtropical zone, the epidemic peaks of HRSV detection were from March to May and from July to August [16–17]. In the temperate zone, the peak of HRSV detection was mainly concentrated in winter and spring from November to April of the following year [18]. Huzhou is located in the temperate zone, HRSV activity peaks in the winter-spring period. Infection with HRSV was found in all age groups tested in Huzhou, the detection rate decreases with the increase of age. The highest detection rate was the 0-1 year age group. These results were consistent with the finding in other places in China [19–21]. It was clear that the most affected population by HRSV infection was children, in particular, children less than one year old.

In China, GA2 was the predominant genotype of HRSV-A from 2003/2004 to 2007/2008[22–25]. However, the GA2 genotype was replaced by the NA1 genotype from 2008 to 2013. The NA1 genotype was first reported in Beijing in 2007 [26] and circulated in other provinces of China in the following years [27–28]. ON1 genotype was first detected in December 2011 in China, and then gradually became the dominant genotype, and steadily replaced the previously circulating NA1 genotype [28–30]. Globally, the ON1 genotype has been reported in many countries and was evolving quickly throughout the world with different ON1 lineages [31]. The HRSV-A strains identified in Huzhou were all ON1 from 2016 to 2019. These results were consistent with the findings in other places in China.

HRSV-B genotype BA was first identified in Buenos Aires, Argentina in 1999. It has a duplication of 60 nucleotides in the second hypervariable region of the G gene [32]. After its emergence in 1999, BA was spreading globally and became the predominant among HRSV-B strains in many countries [33–34]. Since the BA genotype was reported, it has been prevalent as a major genotype in the world for more than 20 years and has evolved into multiple genotypes after these years circulating, indicating that the BA genotype has a strong adaptability [35]. BA9, a branche of BA, was first identified during 2006-2007 season. Then became dominant worldwide from the end of 2009 [36–38]. BA9 genotype was the dominant genotype in Huzhou too. All the HRSV-B viruses found in this study belong to the BA9 genotype.

There is no persistent protective antibody after infection with HRSV. It is common to be infected with HRSV repeatedly. The substitution of amino acids in G protein and the appearance of new glycosylation sites may be the reasons for HRSV to evade immune surveillance and cause repeated infection [39].

There are several limitations in our study. Firstly, we could not sequence all identified HRSV strains, which may have other genotypes. Secondly, we only used partial G protein for genetic analysis. Thirdly, our data covered only the last four years, it may not be enough to draw any conclusion. The ongoing RSV surveillance and the full lengths of G protein should be pursued in future investigations.

## Conclusions

In this study, we analyzed the HRSV strains circulation from January 2016 to December 2019 in Huzhou, China. This is the first molecular analysis on HRSV in Huzhou. We found subgroup A and B of HRSV were co-circulating and the 0-1 year age group having the highest infection rate. Our study data contributes to understanding the HRSV molecular epidemiology in Huzhou.

## List of abbreviations

HRSV: Human respiratory syncytial virus

## Declarations

### Ethics approval and consent to participate

This study was approved by the ethics committee of Huzhou Center for Disease Control and Prevention (20160521). Informed consent for the nasopharyngeal swabs was obtained from the patients or their guardians. This study was part of a routine laboratory-based investigation. No human experimentation was conducted. The only human materials used were nasopharyngeal swabs that had been sent to our laboratory for routine virological diagnosis.

### Consent for publication

Not applicable.

### Availability of data and materials

The readers interested in using the data may contact the corresponding author.

### Authors’ contributions

LPC and DSX participated in the design of the study and performed the statistical analysis. DSX and LJ participated in the HRSV detection. XFW, DSX and WY participated in the genomic amplification for genotyping. WY and LJ participated in the sequence analysis and phylogenetic analysis. LPC drafted the manuscript. All authors read and approved the final manuscript.

## Acknowledgements

We thank the staff of the First People’s Hospital in Huzhou for collecting the samples.

